# Analysis of coding variants in the human FTO gene from the ExAC (gnomAD) Database

**DOI:** 10.1101/2021.03.03.433730

**Authors:** Mauro Lúcio Ferreira Souza, Jaime Viana de Sousa, João Farias Guerreiro

## Abstract

Single nucleotide polymorphisms (SNPs) in the first intron of the FTO gene (alpha-ketoglutarate-dependent dioxygenase) identified by a genome-wide association study (GWAS) in 2007 continue to be the known variants with the greatest effect on adiposity in different human populations. Currently available data reveal a total of 61 different intronic SNPs associated with adiposity. Coding variants in the FTO gene, on the other hand, have been little explored, but data from complete sequencing of the exomes of various populations are available in public databases and provide an excellent opportunity to investigate potential functional variants in FTO. This study aimed to track nonsynonymous variants in the exons of the FTO gene in different population groups using the ExAC database (gnomAD) (http://exac.broadinstitute.org/) and to analyze the potential functional impact of these variants on the FTO protein. Variants were analyzed using five publicly available pathogenicity prediction programs. Of the 158 mutations identified (152 missense and 6 stop-gain), 64 (40.5%) were classified as pathogenic, 67 (42.4%) were classified as benign, and 27 (17%) were classified as inconclusive. Thirty variants were classified as pathogenic by all five predictors used in this study, and 16 mutations were classified as pathogenic by only one predictor. The largest number of mutations was found in Europeans (non-Finnish) (85/158), all with very low frequencies, and half (32/64) of the variants classified as pathogenic by the five predictors used were also found in this population. The data obtained in this analysis show that a large number of rare coding variants classified as pathogenic or potentially pathogenic by different in silico pathogenicity prediction programs are not detected by GWAS due to the low linkage disequilibrium as well as the limitations of GWAS in capturing rare variants present in less than 1.0% of the population.

## Introduction

FTO (alpha-ketoglutarate-dependent dioxygenase) was the first obesity susceptibility gene identified through genome-wide association studies (GWAS) and remains the locus with the greatest effect on adiposity in different human populations. Four independent GWAS published in 2007 reported a significant association between body mass index and body fat and common genetic variants in the FTO gene, specifically, a group of single nucleotide polymorphisms (SNPs) in the first intron of the gene. FTO was identified for the first time in Europeans in 2007 [1], and shortly thereafter, its association with BMI and obesity risk was confirmed by three other studies [2, 3, 4]. This association has been replicated in other populations (Asians, Hispanics and Native Americans), but conflicting results have been observed in African populations [5, 6]. The frequencies of risk alleles vary substantially between different ethnic groups, which may explain, to some degree, the differences in estimates of the effects of these alleles on BMI. Different populations are characterized by several specific patterns of tightly linked SNP haplotypes associated with the phenotype [7]. The FTO gene, located on chromosome 16q12.2, is expressed in a wide range of tissues, as it is a maintenance gene that maintains the CpG islands in gene promoters. It contains nine exons and spans approximately 410 kb, making it unusually large for a maintenance gene. It encodes a 2-oxoglutarate-dependent oxygenase that performs oxidative demethylation of RNA/DNA, and the available data suggest that FTO plays a role in the arcuate nuclei of the hypothalamus, where it mediates energy balance and eating behavior [7]. The intronic location of common SNPs associated with BMI and obesity within a 47 kb region that covers parts of the first two introns and exon 2 of FTO [1] indicates that the amino acid sequence of the FTO protein does not exert its effects through functional mutations and is more likely to play a role in transcription regulation through its impact on the expression of the FTO gene and/or neighboring genes such as the IRX3/IRX5 genes, specifically in adipocytes. Experimental data [8] confirmed that FTO intron 1 is involved in enhancer activation, previously described by another study [9], and regulates the expression of the IRX3 and IRX5 loci, which are vital for the maturation of adipocytes [7, 6].

Data obtained from the NHGRI-EBI GWAS catalog [10] in 2017 revealed a grouping of 15 SNPs associated with obesity in intron 1 of the FTO gene [11], but the data currently available reveal a total of 61 different intronic SNPs associated with BMI, distribution of body fat and other characteristics of obesity, almost all present in Europeans, Africans, East Asians, South Asians and Americans (miscegenated populations in Latin America) and in a lower percentage (19/61) in Native Americans (Peruvian Amerindians) [12]. Coding variants in the FTO gene, on the other hand, have been little explored. However, complete sequencing data for the exome of continental populations are available from public databases such as the Genome Aggregation Database (gnomAD) and the NHLBI Exome Sequencing Project (Exome Variant Server) and provide an excellent opportunity to investigate potentially functional alleles in the FTO gene. Thus, the aim of this study was to track nonsynonymous variants in the exons of the FTO gene in different population groups using the gnomAD database and to analyze the potential functional impact of these variants on the FTO protein.

## Materials and Methods

The FTO gene data available in ExAC (gnomAD) were downloaded from http://exac.broadinstitute.org/. The dataset provides sequence variation in 60,706 unrelated individuals from various population genetic and disease-specific studies across ethnicities, including European (non-Finnish), European (Finnish), African, Latin, South Asia, East Asia and "others". GnomAD, also known as the Genome Aggregation Database Consortium, was developed by an international coalition of researchers to aggregate and harmonize exome and genome sequencing data from a wide range of large-scale sequencing projects and to make data available summaries for the scientific community. Formerly known as the Exome Aggregation Consortium (ExAC), the project started in 2012 and expanded on the work of the 1000 Genomes Project and others that cataloged human genetic variation [13, 14]. The reference genome used for sequence alignment was GRCh37/hg19 (reference), and alignment was performed using the GATK tool [15]. Variants were analyzed using the following publicly available pathogenicity prediction programs: FATHMM [16], PROVEAN [17], SIFT [18], POLYPHEN-2 [19] and PANTHER [20]. The criteria used to classify the nature of the mutations were as follows: benign, when three or more predictors classified the variant as benign; pathogenic, when three or more predictors classified the mutation as pathogenic; inconclusive, when at least one predictor was unable to analyze the variant and two classified it as pathogenic and two others classified it as benign or when no prediction was made by multiple predictors.

ClinVar [21] is one of the most commonly used databases for clinical and pathological analyses of mutations. However, the vast majority of exonic SNPs identified in the FTO gene are not registered in this database and therefore do not have an rsID and could not be verified in this database.

The variant data found in ExAC (gnomAD) were compared with those obtained from the Exome Variant Server (EVS) [22], a database containing variants identified by exome sequencing retrieved from the National Heart Lung and Blood Institute (NHLBI) Exome Sequencing Project (ESP), designed to identify genetic variants in the coding regions (exons) of human genes that are associated with heart, lung and blood diseases. EVS annotation data were generated from approximately 5400 exomes of European and African American populations (https://evs.gs.washington.edu/EVS/).

## Results and Discussion

In total, 158 nonsynonymous mutations were identified in the ExAC database (gnomAD), of which six were stop-gain mutations (3.7%) and 152 were missense mutations (96.3%). Of this total, only 21 variants were found with rsIDs in the Exome Variant Server (EVS) database. Of the 158 mutations identified, 64 (40.5%) were classified as pathogenic, 67 (42.4%) were classified as benign, and 27 (17%) were classified as inconclusive based on *in silico* analyses by five pathogenicity predictors (Table 1).

Information on the position, nucleotide change, amino acid change, type of mutation, allele count, number of alleles and frequency of each variant in Latino, South Asian, East Asian, African, European (non-Finnish) and European (Finnish) populations is shown in Table S1. Of the 38 mutations identified in South Asians, 15 were classified as pathogenic, 12 as benign and 11 as inconclusive. The most frequent mutation (Arg123Trp), classified as inconclusive, was the only one found with a frequency ≥ 1% (2.1%). The other mutations were found at very low frequencies. The most common pathogenic mutation was Glu325Val (0.65%). In East Asian populations, 23 very rare mutations have been identified. Six mutations were classified as pathogenic, 10 as benign and seven as inconclusive. The two most common mutations, Asp144 (0.004) and Asp113 (0.002), were classified as benign and inconclusive, respectively. The variants classified as pathogenic had frequencies in the range of 0.0001. Thirty-one mutations with very low frequencies, in the range of 1/10,000 or more, have been found in African populations. Of these mutations, eight were classified as pathogenic, 13 as benign and 10 as inconclusive. In Europeans (non-Finns), a greater number of mutations was found (85), but all with very low frequencies, in the range of 1/10,000 or more. Of these, 32 were classified as pathogenic, 35 as benign and 18 as inconclusive. In Europeans (Finns), on the other hand, only eight variants were found. Five were classified as benign, and three were classified as inconclusive. No pathogenic mutations were found. The most common variant, Asp332Gly, classified as inconclusive, had a frequency of 0.004. The others had frequencies in the range of 0.0001. A total of 24 rare mutations have been identified in Latino populations. Of these, 13 were classified as pathogenic, five as benign and six as inconclusive. The most common variant in Latino populations was Arg84Ser (0.002), a variant classified as pathogenic. Globally, the most common variant, Arg123Trp, was found in 296 individuals (0.003); this variant was most common in South Asians (0.021) and was also found in East Asians, Europeans (Non-Finns), Africans and Latinos. However, the pathogenicity of this variant was classified as inconclusive by all five predictors used in this study.

Among the 64 variants classified as pathogenic, half were found in non-Finnish Europeans (32). In South Asians, 15 mutations were identified: in Latinos 13, in Africans eight, and in East Asians six, while in Europeans (Finns), no pathogenic mutations were identified. Globally, the most common pathogenic variant was the Glu325Val substitution, found with a frequency of 0.006 in South Asians and in Europeans (non-Finns) and in Latinos, with very low frequencies. Eleven pathogenic mutations are shared by more than one population group, seven of which are found in Europeans and in one or two other continental populations, suggesting a European origin for the variant with spread by migration: the variants Tyr333Cys, Pro399Ala and His62Arg for Latinos, a Glu325Val variant for Latinos and South Asians, the Arg322Gln and Arg388Ter variants for South Asians, and the Arg337Cys variant for East Asians.

Data on the pathogenicity of variants according to the five prediction programs used in this study are shown in Table S2. As expected, there were conflicts between the results of the pathogenicity analysis of the variants performed by the five prediction programs used in this study. In general, the FATHMM and PROVEAN programs classified most variants as benign (90; 57% and 80; 50.6%, respectively), while the programs SIFT, POLYPHEN and PANTHER classified most variants as pathogenic (70; 44.3%, 94; 59.5% and 80; 50.6%, respectively (Table 3). Pathogenicity prediction programs allow the evaluation of the effect of amino acid substitutions on the structure or function of a protein without performing functional studies, and the available data show that the average accuracy of pathogenicity predictors is 85%. However, as the different pathogenicity prediction programs vary widely in their methods and ability to predict the pathogenicity of a given sequence change, there are significant disagreements in the identification of mutational impact and pathogenicity among the different programs [23, 24]. In total, 30 exonic variants were classified as pathogenic by all five predictors used in this study, and 16 variants were classified as pathogenic by at least one of the pathogenicity prediction programs (Table S2).

The comparison of the FTO exonic variant data identified in ExAC was performed with data available in the Exome Variant Server database obtained from European and African-American populations, which uses only the Polyphen2 program as a pathogenicity predictor. The analysis revealed a total of 21 variants shared between ExAC and EVS. In total, in this study, 40.5% of the ExAC variants were classified as pathogenic, whereas 25.3% were classified as probably/possibly pathogenic in the EVS. The proportions of variants classified as benign were 42.4% and 52.5% in this study and in the EVS, respectively, while the proportions of variants classified as inconclusive in this study and in the EVS were 17.1% and 52%, respectively (Table 1). Four variants (Asp332Gly, Arg337Cys, Arg455Ser and Leu489Phe) of ExAC were classified as pathogenic in this study and probably/possibly pathogenic in EVS, six were classified as benign in this study and probably/possibly pathogenic in EVS, and nine were classified as benign in this study and in the EVS (Table 2). Exonic variants of the FTO gene have been little explored, and the few studies to screen for variants by exon sequencing of the FTO gene have found no evidence that the identified variants confer an increased risk of obesity. In obese European children (English, French, Belgian and Swiss), 34 variants were identified [25], while only seven nonsynonymous variants were found in Chinese (Han) children with early-onset obesity [26]. Likewise, next-generation sequencing (NGS) of the FTO gene in severely obese Swedish children has revealed little evidence of functional variants in the coding region of the FTO gene [27]. However, complete exome sequencing data for large populations available from public databases such as The Genome Aggregation Database (gnomAD) and NHLBI Exome Sequencing Project (Exome Variant Server) provide an excellent opportunity to identify potential functional variants in the FTO gene in a larger number of populations globally. The data obtained in this analysis show a substantial number of rare coding variants classified as pathogenic or potentially pathogenic by different pathogenicity prediction programs, which are not detected by GWAS due to the low linkage imbalance as well as the limitations of GWAS in capturing rare variants present in less than 1.0% of the population. Therefore, this type of analysis can be an adequate alternative to selecting target SNPs for further exploration through confirmatory functional tests or in case-control studies and twin studies through next-generation sequencing platforms.

## Acknowledgments

We thank Dr. Gilderlânio Santana de Araújo for valuable comments and suggestions on the manuscript.

